# Integrating tDNA Epigenomics and Expression with Codon Usage Unravel an Intricate Connection with Protein Expression Dynamics in *Trypanosoma cruzi*

**DOI:** 10.1101/2024.07.04.602108

**Authors:** Herbert G. S. Silva, Satoshi Kimura, Pedro L. C. Lima, David S. Pires, Matthew K. Waldor, Julia P. C. da Cunha

## Abstract

Codon usage bias impacts protein expression across all kingdoms of life, including trypanosomatids. These protozoa, such as the *Trypanosoma cruzi*, primarily regulate their protein-coding genes through posttranscriptional mechanisms. Here, we integrated analyses of codon usage with multiple high-throughput sequencing data to investigate whether codon usage bias is present into surface virulence factors (disruptive compartment), conserved housekeeping proteins (core compartment), and proteins involved in the developmental stages of *T. cruzi*. For the first time in trypanosomatids, tRNA sequencing was employed to reveal coadaptation between codon usage and anticodon availability. Despite notable differences in the proteomes of infective and non-infective forms, they exhibited similar pools of tRNAs and similar codon usage preferences. We observed that open chromatin levels of tRNA genes correlate with tRNA expression in non-infective forms, but not in infective forms, suggesting chromatin states do not control the tRNA pool in the latter. Our analysis also revealed a relationship between anticodon:codon pairing modes and protein abundance. Highly expressed mRNAs favored Watson– Crick base pairing, whereas less expressed mRNAs displayed more wobble base pairing. Overall, our findings suggest that protein expression in *T. cruzi* is influenced by a combination of codon usage bias, tRNA abundance, and anticodon:codon pairing modes.

**Graphical abstract:** 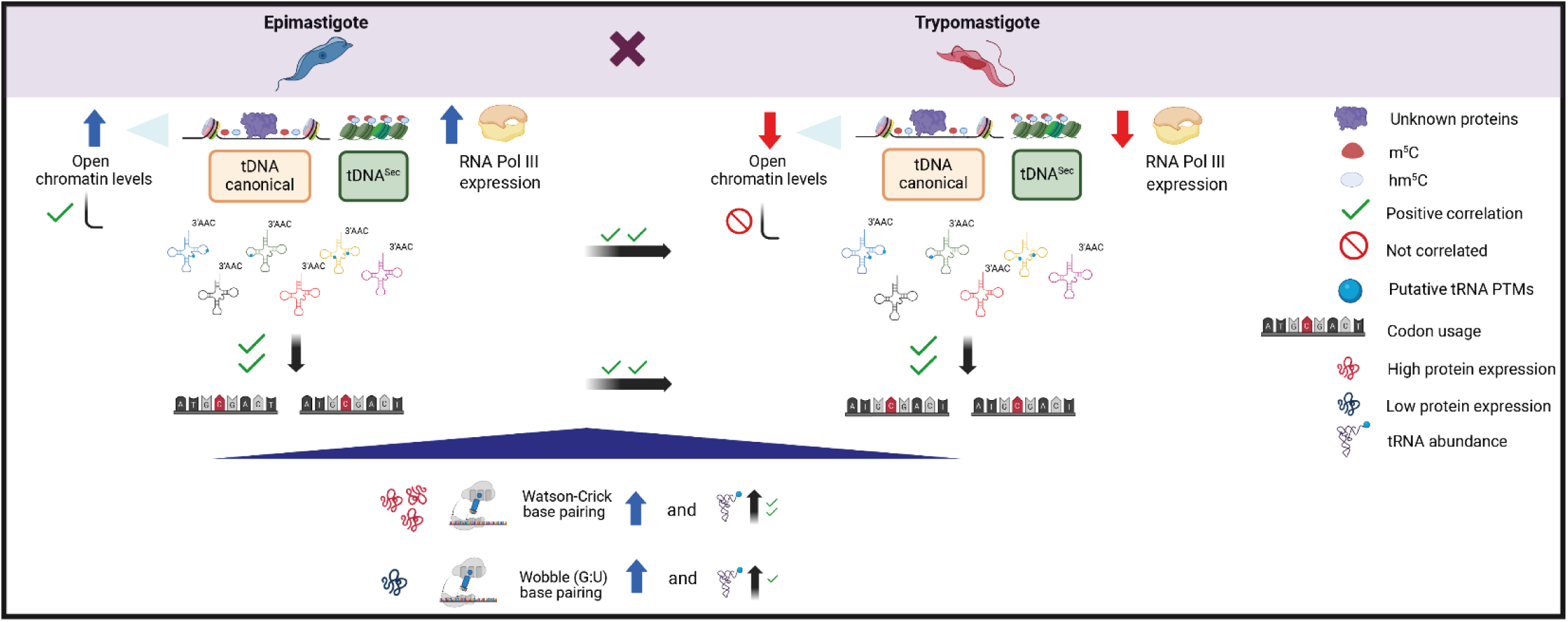

## Introduction

The genetic code, defined by triplet codons in messenger RNAs (mRNAs), dictates the amino acid sequence of proteins. Typically, sixty-four codons are present in the genome of most organisms, each encoding one of the twenty common amino acids, except for three stop codons (UGA, UAA, UAG) involved in translation termination (Crick, 1968). Most amino acids are encoded by two to six ‘synonymous’ codons (Crick, 1968). The biased usage of synonymous codons, where a subset of genes has a different codon bias (nonuniform usage of synonymous codons) from that of the remainder of the genome, has been observed among different organisms (Bahiri-Elitzur and Tuller, 2021). Such synonymous codon usage bias has been reported to contribute to the optimization of translation and mRNA stability (Bahiri-Elitzur and Tuller, 2021; Gingold et al., 2012; Hershberg and Petrov, 2009; Jeacock et al., 2018; Liu, 2020; Rudolph et al., 2016; Sharp and Li, 1987). However, our understanding of the impacts of codon usage bias on translation remains limited in many organisms.

Synonymous codon usage bias is linked to the availability of transfer RNA (tRNA) (Bahiri-Elitzur and Tuller, 2021; Hanson and Coller, 2018; Rak et al., 2021; Rudolph et al., 2016; Sabi and Tuller, 2014). tRNAs, which are linked to specific amino acids, have distinct anticodon sequences that bind to the triplet codons in mRNA, thereby translating the genetic code into amino acids in the A site of ribosomes (J D WATSON, 1964; Warner and Rich, 1964). The correspondences between tRNA anticodons and mRNA codons are complex. The pairing of the anticodon-codon sequences could be mediated by different interactions, such as Watson–Crick and wobble base pairing. The Watson‒Crick interaction demands a precise match of the triplet, following the strict rules of complementary pairing (Cleaves, 2011; WATSON and CRICK, 1953). On the other hand, wobble base pairing via inosine modification or G:U interactions allows flexibility at the first position of the anticodon (5’-3’) to recognize some ribonucleotides at the third position of the codon (5’-3’). For instance, inosine bases can pair with A, U or C in codon sequences, while G pairs with U in the anticodon:codon interaction context (Agris et al., 2018; Crick, 1966; Varani and McClain, 2000). Thus, in many cases, a tRNA binds to multiple synonymous codons, and a single codon can be decoded by multiple tRNA species. Such redundancy in codon:anticodon pairing enables balanced and robust codon decoding.

The balance between codon usage patterns and the availability of tRNAs is crucial for efficient translation (Bahiri-Elitzur and Tuller, 2021; Hanson and Coller, 2018; Rak et al., 2021; Rudolph et al., 2016; Sabi and Tuller, 2014). Matching of the supply of tRNAs to the demand for codons is necessary for rapid and constant rates of translation. A low availability of tRNA leads to reduced rates of translation and ribosome pausing, which leads to reduced protein production and sometimes triggers stress responses (Kato et al., 1990; Liu, 2020).

tRNA abundance is regulated through genetic, epigenetic, and posttranscriptional mechanisms, including tRNA modifications (Cui et al., 2023; Mahapatra et al., 2011; Ohira and Suzuki, 2024; Shukla and Bhargava, 2018; Yared et al., 2024). It has been reported that tRNA pools can be adjusted in response to different environmental cues, enabling rapid and increased synthesis of proteins essential for cellular adaptation to environmental changes (Torrent et al., 2018). Substantial changes in the expression of different tRNA isodecoders are observed in different mouse tissues (Pinkard et al., 2020) and during the differentiation of human cells (Gao et al., 2024), although the overall tRNA anticodon pools (related to the decoding rates) are largely stable. Therefore, profiling the dynamics of tRNA abundance is critical for understanding the impact of codon usage bias on protein synthesis.

Several methods have been used to estimate tRNA abundance profiles. tRNA genomic copy numbers have been widely used since, in many eukaryotes, they correlate with the abundance of their corresponding codons (Du et al., 2017; Iben and Maraia, 2012; Liu et al., 2021). Although they are widely used, it is not clear whether they reflect the actual tRNA expression profiles in most organisms or under different conditions. The recent development of tRNA sequencing has enabled direct profiling of tRNA expression, but this technique has been applied only to a limited number of organisms and under a limited number of conditions. Direct profiling of tRNA abundance and its correlation with codon usage bias is crucial for understanding the principles of global translation regulation.

Trypanosomatids are a group of unicellular parasitic protozoa that include pathogens such as *Trypanosoma cruzi* (*T. cruzi*), *Trypanosoma brucei* (*T. brucei*) and *Leishmania major* (*L. major*). These protozoa exhibit polycistronic transcription of mRNA genes, and their gene expression regulation occurs mainly posttranscriptionally (Clayton, 2019). *T. cruzi* is the etiological agent of Chagas disease and has four different developmental stages: two infective and nonproliferative stages (metacyclic trypomastigotes - MTs and tissue culture trypomastigotes - TCTs) and two noninfective and proliferative stages (epimastigotes - EPIs and amastigotes) (de Souza et al., 2010). *T. cruzi* differentiation from EPIs to MTs is followed by changes in gene expression, which is important for the parasite’s virulence and survival in its new habitat (de Godoy et al., 2012; Freire-de-Lima et al., 2015; Smircich et al., 2015).

The *T. cruzi* genome is organized into conserved syntenic segments within trypanosomatids (the core compartment) and a nonsyntenic segment (the disruptive compartment) composed of virulence factor genes associated with multigene families, such as trans-sialidase (TS), mucins, and maspins (MASP) (Berná et al., 2018). GP63, dispersed gene family 1 (DGF-1), and retrotransposon hot spot (RHS) multigene families can be found in either compartment (Berná et al., 2018); when one of these families is found in a given compartment, the compartment is referred to as the GpDR compartment. EPIs predominantly translate genes from the core compartment, while MTs have increased production of disruptive proteins (de Godoy et al., 2012; Smircich et al., 2015). These distinct compartments have different GC contents (Berná et al., 2018), suggesting that distinct codon usage biases are likely associated with parasite development. Codon bias has been proposed as an important mechanism to direct global relative mRNA and protein expression levels in trypanosomatids (de Freitas Nascimento et al., 2018; Jeacock et al., 2018). However, a poor correlation between codon usage frequency and the copy number of tRNA genes (tDNAs) was observed in *T. cruzi* (Padilla-Mejía et al., 2009), suggesting that tRNA abundance is controlled by additional mechanism(s). However, how codon usage and tRNA profiles affect the *T. cruzi* translatome and development is largely unclear.

Here, we combined multiple genome-wide datasets to investigate the relationships among tRNA abundance, codon usage, and gene expression profiles in *T. cruzi*. Codon usage analysis revealed that the two *T. cruzi* genomic compartments, as well as transcripts of higher and lower abundance, exhibit codon bias. We adopted tRNA sequencing in two developmental forms (EPI and MT) to profile tRNA abundance. Despite the distinct tRNA chromatin states in these two forms, their tRNA profiles were very similar, suggesting that another mechanism controls tRNA levels across different developmental forms. We observed that highly translated genes predominantly use Watson–Crick base pairing, while poorly translated genes rely more on wobble pairing. This differential use of anticodon:codon pairing modes, together with tRNA availability, may represent a previously undescribed layer of gene expression regulation in *T. cruzi*.

## Materials and methods

### Cell culture

*T. cruzi* (Dm28c strain) EPIs were cultivated at 28 °C in Liver Infusion Tryptose (LIT) medium supplemented with 10% fetal bovine serum (FBS; Vitrocell), 0.4% glucose, 0.1 µM hemin, and 60 mg/mL penicillin G, as described by Camargo *et al*, 1964. The metacyclic trypomastigote forms were obtained following the protocol outlined by Contreras et al., 1985 (Contreras et al., 1985) with some modifications. In brief, epimastigotes in the exponential growth phase (4×10**^6^** parasites/mL) were cultured for 4 days until they reached the stationary phase (5-6×10**^7^** parasites/mL). Subsequently, the parasites were resuspended at a concentration of 5×10**^8^** parasites/mL in Triatomine Artificial Urine (TAU) medium (190 mM NaCl, 17 mM KCl, 2 mM CaCl_2_, 2 mM MgCl_2_, and 8 mM phosphate buffer, pH 6.0) and incubated for 2 hours at 28 °C. Then, the parasites were diluted to 5×10**^8^**/mL in TAU 3AAG medium (containing 10 mM L-proline, 50 mM L-glutamate, 2 mM L-aspartate, and 10 mM glucose) and maintained in a CO_2_ incubator at 28 °C. Metacyclic forms of the *T. cruzi* Dm28c strain were harvested from the culture supernatant after 76 hours of incubation and purified using DEAE-cellulose resin (Sc-211213).

### tRNA species identification

The *T. cruzi* Dm28c (version 42-2018) genome and its coordinates were downloaded in FASTA and gff formats, respectively, from the TritrypDB platform (https://tritrypdb.org/tritrypdb/app/downloads). To obtain the list of tDNAs and their respective coordinates, the term “tRNA” was used as a filter in the gff file. Identification of selenocysteine tRNA (tRNA^Sec^) was performed using the BLASTp tool v.2.10.0 (Altschul et al., 1990) to search for the best hit via the TriTrypDB platform (https://tritrypdb.org/tritrypdb/app/workspace/blast/new). The query reference was “tRNA selenocysteine” (ID=Tb927.9.2380) from *T. brucei* TREU927. tRNAscan-SE software (http://lowelab.ucsc.edu/tRNAscan-SE/) (Chan et al., 2021) was used to identify the transcribed sense strands and anticodons of the tDNAs. Afterward, a genome FASTA file containing all identified tDNA sequences was obtained.

### tRNA sequencing

Total RNA was extracted using TRIzol (Invitrogen) following the manufacturer’s instructions. tRNA purification and sequencing were performed according to Kimura *et al*., 2020 (Kimura et al., 2020) with some modifications. Five micrograms of RNA samples were run on a 10% denaturing gel (TBE-UREA), and bands corresponding to tRNAs (70 to 85 bp) were excised, digested, eluted, and recovered by isopropanol precipitation. Subsequently, 250 ng of purified tRNA was deacetylated with 500 µL of 100 mM Tris-HCl (pH 9.0) solution and incubated for 1 hour at 37 °C. The tRNA was recovered using isopropanol. Afterward, library preparation for sequencing was carried out as described by Kimura et al., 2020. The tRNA samples were sequenced in biological duplicates for EPIs and MTs on the Illumina NextSeq 1000 system (single-end). The removal of adapters (AGATCGGAAG) from the FASTQ files was performed using cutadapt v.3.5 (Martin, 2011), followed by HEADCROP:2 using the TrimmomaticSE v.0.39 tool (Bolger et al., 2014). The tDNA FASTA file generated in this study (see Supplementary Table 2A) was used for mapping. The **i.** mapping, **ii.** abundance counting, **iii.** Analysis of differential tRNA expression, counting of 3’CCA on tRNA and misincorporation frequencies between epimastigote and trypomastigote metacyclic forms were performed using the mim-tRNAseq script with the parameters --cluster-id 0.97, --max-mismatches 0.075, --max-multi 4 and --max-multi 4 (Behrens et al., 2021), executed in Python 3.9.18 and R 4.3.1. The tRNA-seq dataset were deposited at PRJNA1124437.

### Identification of RNA polymerase III subunits in the *T. cruzi* Dm28c genome

The sequences of the RNA polymerase III subunits (RPC1, RPC2, RPC40, RPC19, RPB6, RPB5, RPB8, RPB10, RPC10, RPC11, RPC17, RPC25, RPC82, RPC53, RPC37, RPC34, and RPC31) of *Saccharomyces cerevisiae* S288c were downloaded from the FungiDB database (https://fungidb.org/fungidb/app). Subsequently, two approaches were employed to identify orthologs in the *T. cruzi* Dm28c strain (version 42–2018) genome. The first approach involved using the BLASTp tool v.2.10.0 (Altschul et al., 1990), which is available on the TriTrypDB platform (https://tritrypdb.org/tritrypdb/app/) to search for alignments with an E-value ≤0.001, followed by confirmation of the protein domain using the platform (https://www.ebi.ac.uk/interpro/search/sequence/). For the unidentified RPC31 protein homologous using the BLASTp method, probabilistic models were created with the HMMER Algorithm v.3.3 (Mistry et al., 2013) based on orthologous sequences of RPC31 from several eukaryotes. Subsequently, a search for the best hits was conducted within the *T. cruzi* Dm28c strain (version 42–2018) genome using the Ubuntu v.20.04 system.

### FAIRE-seq, MNAse-seq, modified bases analysis in EPI and TCT forms

The RPGC values available from FAIRE-seq data performed in the EPI and MT forms were obtained from Lima et al, 2021(Lima et al., 2022). Raw files from MNase-seq data in the EPI and TCT forms were obtained from the SRA project number PRJNA665060 (Lima et al., 2021). The reads were processed using Trimmomatic (Bolger et al., 2014) to remove adapter sequences, and the following parameters were applied: HEADCROP:5, SLIDINGWINDOW:15:25, and MINLEN:35. The rRNA reads were masked using bedtools maskfasta v2.31.1. Reads mapping was performed using Bowtie2 (Langmead and Salzberg, 2012) to the *T. cruzi* Dm28c genome (version 42–2018) available on TriTrypDB using the following parameters: -D 25, -R 4, -N 0, -L 19, and -i S,1,0.40. SAM files were converted to BAM files, which were subsequently sorted and indexed using SAMtools version 1.12 (Li et al., 2009). Read count values for tDNA regions, selenocysteine loci, and shuffled regions equivalent to tRNAs (generated by running Bedtools v2.31.1 with the shuffle option) were obtained using BAMscale Version v1.0 (Pongor et al., 2020). The resulting counts in FPKM were analyzed using libraries such as readr, dplyr, reshape2, ggpubr, and ggplot2 in R Version 4.1.2. Modified bases analysis was performed as described in Lima *et al*, 2024 (Lima et al., 2024). In short, public Oxford Nanopore sequencing datasets available at Díaz-Viraqué *et al*, 2023 (Díaz-Viraqué et al., 2023) were converted into POD5 files using pod5 library version 0.3.6 (pod5 convert fast5), which were used to call methylation (m) and hydroxymethylation (h) cytosines with the Dorado program, version 0.5.3, followed by read mapping using the minimap2 program (Li, 2018) against the genome of the Dm28c 2018 strain (version 60). The analysis of methylated bases was performed using the Modkit program, version 0.2.5 (available at https://github.com/nanoporetech/modkit).

### Ribo-seq data analysis of the *T. cruzi* Dm28c genome

Ribo-seq data for EPI and MT forms of *T. cruzi* were obtained from the Sequence Read Archive (RCA) repository, project ID: PRJNA260933, provided by (Smircich et al., 2015). The FASTQ files were filtered using the TrimmomaticSE v.0.39 tool (Bolger et al., 2014) with parameters –HEADCROP:1 and MAXINFO:24:0:0.05 and mapped to the *T. cruzi* Dm28c (version 42–2018) genome using Kallisto’s algorithm to obtain the transcripts per million (TPM) values with the following settings: -b 100–l 50–s 20 (Bray et al., 2016). The DESeq tool v.1.42 (Love et al., 2014) was used to perform differential expression analysis of the translatome between the EPI and MT forms.

### Codon usage analysis in the *T. cruzi* Dm28c genome

The aa.usage.py script available at https://github.com/rhondene/Codon-Usage-in-Python was used to compute the **i.** relative synonymous codon usage (RSCU), **ii.** codon relative adaptive weights, **iii**. The absolute codon frequencies and amino acid frequencies are shown in FASTA files. The identified genes for the core, disruptive and GpDR compartments were obtained from Lima *et al*, 2021 (Lima et al., 2022). All upregulated CDSs between EPIs (1,752 genes) and MTs (361 genes) were generated from Ribo-seq data, excluding pseudogenes. The top 100 and bottom 100 genes (reference set) were obtained from the Ribo-seq data containing the CDSs with 100 genes with higher or lower TPM values in the EPI and MT forms. The reference set was selected based on genes with a log2-fold change of ≤ 2 (EPIs vs. MTs) and an adjusted p value of ≤ 0.05, excluding pseudogenes and genes present in multiple copies. The tRNA abundances, represented as the proportion of mapped reads, were determined for EPIs and MTs. To refine these measurements, a renormalization process was applied to address the potential for a single tRNA to recognize multiple codons. This was achieved by dividing the abundance of each tRNA by the number of its corresponding codons and then assigning these adjusted abundance values to each codon. For codons recognized by multiple tRNA species, the total abundance was calculated by summing the adjusted abundances of all tRNA species recognizing that codon.

## Results

### Core and disruptive genomic regions have differences in codon usage

The core and disruptive genomic compartments differ in their GC content, gene class diversity, evolutionary history, and their protein-coding genes are differentially expressed in *T. cruzi* life forms (Berná et al., 2018). EPIs predominantly translate core genes, while MTs exhibit elevated expression of disruptive transcripts (de Godoy et al., 2012; Smircich et al., 2015). This pronounced shift in their respective proteomes raised the question of whether these two genomic compartments, core and disruptive, display different patterns of relative frequencies of amino acids and codons in their coding regions. We observed that the codon frequencies of the core genes were different from those of the disruptive and GpDR genes (Figure 1A). Genes from the core and disruptive regions had high correlation levels when amino acid (R=0.81) or codon absolute frequencies (R=0.75) were analyzed (Figure 1B-C and Supplementary Table 1A). Ala and Gly had the highest amino acid frequencies for the core and disruptive compartments, respectively. Codons for Glu were among the highest for both compartments.

**Figure 1.**
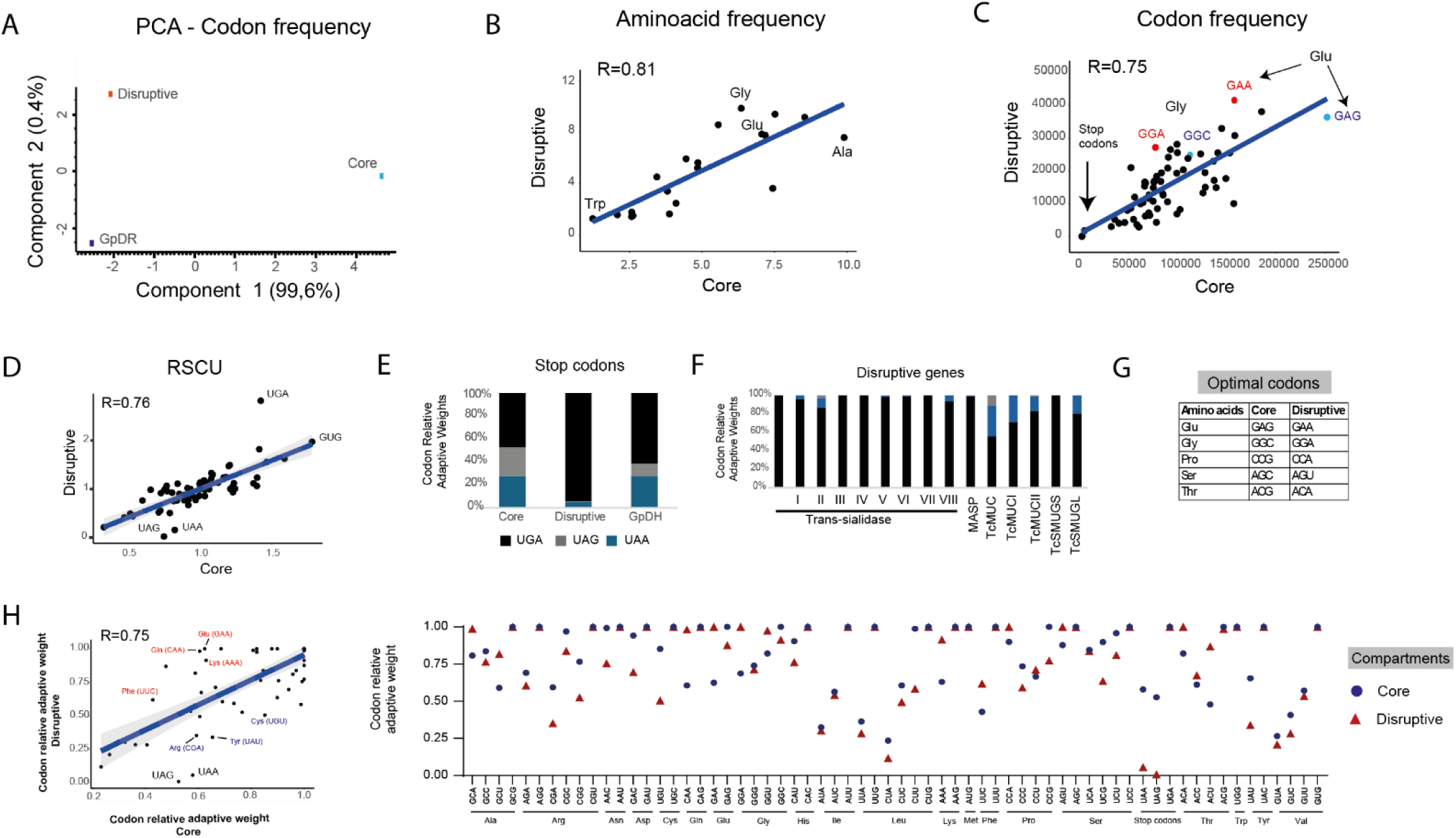
Identification of codon usage bias in the *T. cruzi* genome. **A)** Principal component analysis (PCA) of codon absolute frequencies for the core, disruptive and GpDR compartments. Scatter plot for either **B)** amino acid or **C)** codon usage frequencies in core and disruptive nucleotide sequences. Pearson’s correlations are shown above each graph. **D)** Scatter plot based on Pearson’s correlation of RSCU values for core and disruptive genomic compartments. **E)** Percentage of codon relative adaptive weights for stop codons (UGA, UAG and UAA) in the core, disruptive and GpDR compartments. **F)** Enrichment of the stop codons on the class of disruptive genes: trans-sialidases, MASP, and mucins. **G)** Table displaying the distinct optimal codons between the core and disruptive compartments. **H)** Relative adaptive weights of codons compared to those of optimal codons from core and disruptive compartments. The highlighted codons are at least 20% different between the indicated conditions. The Pearson’s correlation between both compartments is depicted on the left side, with the respective values for each codon presented on the right side. Gray shading, 95% confidence interval.

To obtain further insights into codon bias, we compared the relative synonymous codon usage (RSCU) values for core and disruptive CDSs. RSCU indicates the preference for specific codons that encode the same amino acid or translation termination codon (Li et al., 2023; Sharp and Li, 1987). Higher RSCU values indicate optimal codons, which are associated with more abundant corresponding tRNA species for those amino acid-encoding codons (Hanson and Coller, 2018). The GUG and UGA codons exhibited the highest RSCU values for the core and disruptive CDSs, respectively (Figure 1D). Interestingly, UGA is one of the three stop codons, and in accordance, the relative codon adaptive weights for stop codons differ greatly among genomic compartments. Strikingly, UGA stop codons represented 94% of stop codons in disruptive genes but only 47% and 61% of stop codons in the core and GpDR compartments, respectively (Figure 1E). Among disruptive genes, we detected that TS and MASP genes use UGA as a stop codon for almost 100% of their genes (Figure 1F). In addition, we detected five distinct optimal codons for the amino acids Glu, Gly, Pro, Ser and Thr in the core and disruptive genes (Figure 1G and Supplementary Table 1A). Moreover, when analyzing codon relative adaptive weights compared to their respective optimal counterparts, an evident bias for Arg (CGA), Cys (UGU), and Tyr (UAU) was found for the core CDSs, while Gln (CAA), Glu (GAA), Lys (AAA) and Phe (UUC) were more prevalent in the disruptive CDSs (Figure 1H and Supplementary Table 1A). Collectively, although the core and disruptive genomic regions have similar frequency of amino acid and codon usage (Figure 1 B and C), these two genetic regions demonstrated different preferences in synonymous codon usage and optimal codons.

### Characterization of tRNA isotypes and anticodon:codon pairing modes

In the *T. cruzi* hybrid strain CL Brener, the genome contains forty-six anticodons, allowing for the translation of sixty-two codons representing the 20 canonical amino acids and one noncanonical amino acid, selenocysteine (Sec) (Padilla-Mejía et al., 2009). The nonhybrid *T cruzi* strain Dm28c displays a lower tDNA copy number, with a total of 105 tDNAs compared to CL Brener, which has 120 copies (Padilla-Mejía et al., 2009; Viraqué et al., n.d.). Our study focused on the *T. cruzi* Dm28c strain, which, similar to the CL Brener strain, has poorly characterized anticodon:codon pairing modes. Thus, we used the tRNAscan-SE tool to characterize the isoacceptors and isodecoders and subsequently established their respective base pairings with the sixty-two codons present in the *T. cruzi* Dm28c genome. We identified forty-six isoacceptors (anticodons) and eight isodecoders in the *T. cruzi* Dm28c strain (Figure 2A) that specifically carry one of twenty canonical amino acids found in nature, as well as one tRNA that carries the noncanonical amino acid selenocysteine. The majority of tRNA species are present as a single- or two-gene copies, as depicted in Figure 2A.

**Figure 2.**
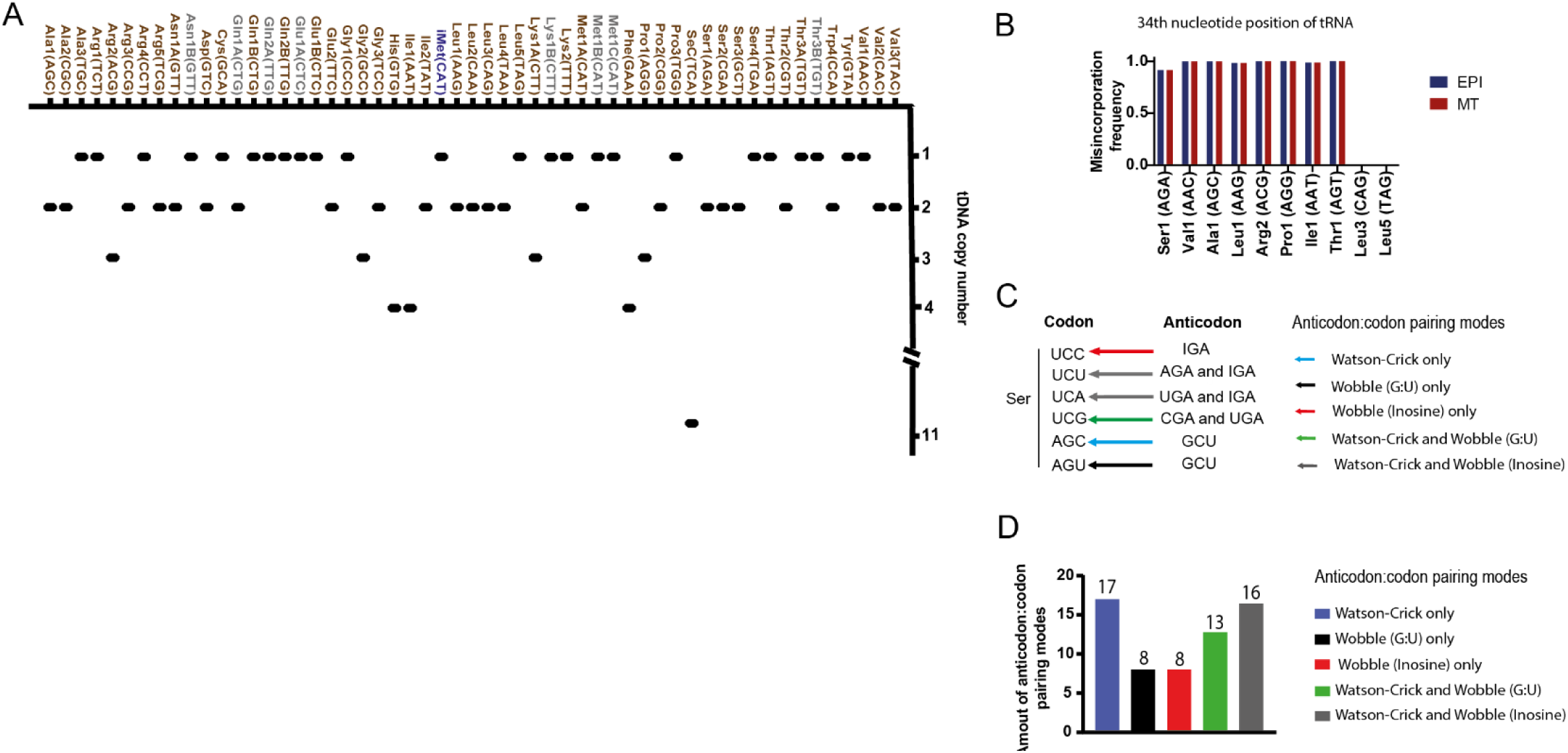
Identification of tRNA isotypes and anticodon:codon base pairing modes in the *T. cruzi* (Dm28c) genome. **A)** tDNA copy number for all tRNA species in the *T. cruzi* genome. The tRNA isoacceptors and isodecoders are represented by brown and gray, respectively. tRNA^imet(CAT)^ is represented in blue. **B)** RT signature (misincorporation frequency) for inosine modification at the 34 nucleotide position of tRNAs in the EPI and MT forms obtained from tRNA-seq data. tRNA^Leu3 (CAG)^ and tRNA^Leu5 (TAG)^ were used as negative controls for the absence of this modification. **C)** Illustration of the anticodon:codon base pairing modes found in the *T. cruzi* genome. **D)** Number of anticodon:codon pairing modes (Watson–Crick only, **ii.** wobble (G:U) only, **iii.** wobble (inosine) only, **iv.** Watson–Crick and wobble (G:U) and **v.** Watson–Crick and wobble (Inosine)).

In addition, we evaluated whether tRNA anticodons containing adenosine at the first position (5’-3’) could undergo modification to inosine (I) in the EPI and MT forms as a strategy to complement codons with missing anticodon matches. The I modification provides wobble base pairing flexibility between synonymous codons. For instance, tRNA^Ala(CGA)^ modified to tRNA^Ala(CGI)^ recognizes the codons ending (5’-3’) with U, C and A as Ala (GC**U**), Ala (GC**C**), and Ala (GC**A**), expanding its recognition beyond Ala (GCU). The detection of I can be achieved using tRNA-seq data, as this modification induces high levels of nucleotide misincorporation (90% to 100%) at modified base positions during reverse transcriptase (RT) from RNA to cDNA, thereby generating an RT signature (Kimura et al., 2020; Schwartz and Motorin, 2017). Thus, we sequenced the total tRNAs from both the EPI and MT forms in duplicate, and the reads were mapped onto a FASTA file genome composed of fifty-five tRNA sequences (Figure Supplementary S1A). The sequencing reads exhibited high quality, coverage and reproducibility between the duplicates (Figure Supplementary S1B-D). We found that all tRNA anticodons beginning (5’-3’) with adenosine in *T. cruzi* are modified by inosine, including tRNA^Ser1 (AGA)^, tRNA^Val1 (AAC)^, tRNA^Ala1 (AGC)^, tRNA ^Leu1 (AAG)^, tRNA^Arg2 (ACG)^, tRNA^Pro1 (AGG)^, tRNA^Ile1 (AAT)^, and tRNA^Thr1 (AGT)^. These modifications remain stable when comparing the EPI and MT forms; they had no detectable changes in their levels across both life forms (Figure 2B).

Based on the variations of anticodons in tRNA genes, the all anticodon:codon base pairing possibilities in *T. cruzi* (**i.** Watson–Crick only, **ii.** wobble (G:U) only, **iii.** wobble (inosine) only, **iv.** Watson–Crick and wobble (G:U) and **v.** Watson–Crick and wobble (Inosine)) are illustrated in Figure 2C, in which seventeen codons display only Watson–Crick pairing with their corresponding tRNA anticodon, eight display either wobble (G:U) or wobble (inosine) pairing, thirteen display Watson–Crick and wobble (G:U) pairing and sixteen codons are recognized by Watson–Crick and wobble (inosine) (Figure 2D and Supplementary Table 2A and B).

### The relative abundance of tRNAs is similar in the EPI and MT forms, despite differences in their posttrancriptional modifications

The global quantification of tRNA species through sequencing has significant challenges, primarily due to the presence of numerous nucleotide modifications in tRNA molecules. These modifications can lead to either nucleotide misincorporation or incomplete conversion of RNA to cDNA during library construction, leading to challenges in mapping and quantification (Behrens et al., 2021). Additionally, the presence of multiple isodecoders, which contain variations in tRNA sequences outside of the anticodon, makes accurate mapping difficult. To circumvent these challenges, we used a highly processive reverse transcriptase for efficient library construction and the mim-tRNAseq analysis pipeline (Behrens et al., 2021; Kimura et al., 2020). The EPI and MT forms of *T. cruzi* displayed a very strong positive correlation (Pearson’s *R* =0.91) related to their tRNA abundance (Figure 3A). Three (**i**.e., Asn1A (GTT)/Asn1B (GTT); **ii**. Lys1A (CTT)/Lys1B (CTT); and **iii**. Gln2A (TTG)/Gln2B (TTG) (Figure 3B and Supplementary Table 2C) of the eight isodecoders could not be definitively mapped to their isoacceptors due to the high similarity between these sequences and the lack of unique mapped reads.

**Figure 3.**
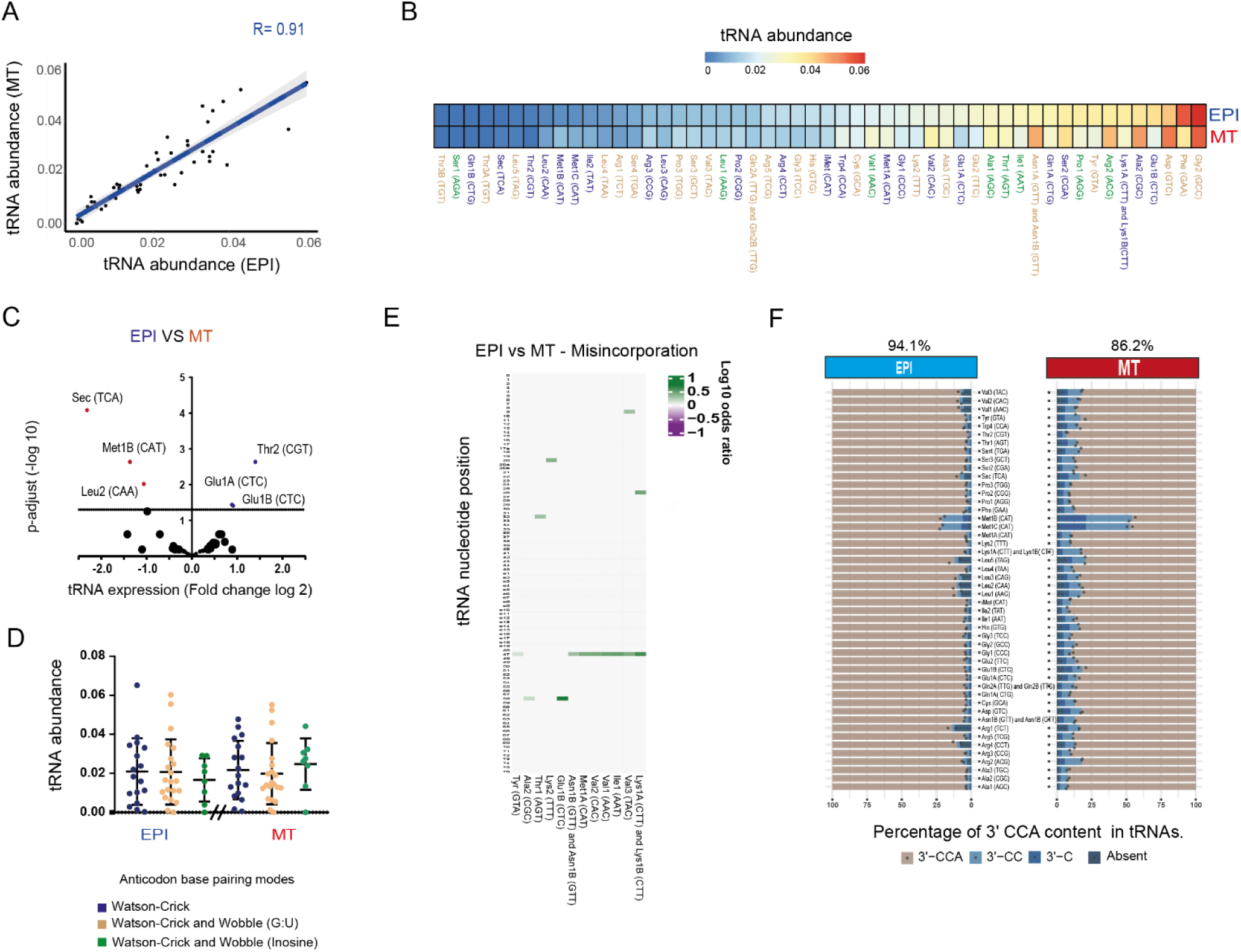
Abundance of tRNA transcripts in EPIs and MTs. **A)** Scatter plot showing Pearson’s correlation between tRNA abundance (proportion of mapped reads) in the EPI and MT forms. Gray shading, 95% confidence interval. **B)** Heatmap with tRNA abundance levels normalized by the proportion of mapped reads on the tDNA genome from *T. cruzi*. tRNA anticodons that exhibit only Watson–Crick base pairing with synonymous codons are highlighted in blue. The tRNA anticodons containing either Watson–Crick or wobble base pairing are represented by orange, while those interactions of either Watson–Crick or inosine are illustrated in green. **C)** Volcano plot of fold changes of tRNA expression (Log2) in EPIs compared to MTs. The significance of tRNA differential expression was determined using the DESeq2 algorithm considering a p-adjust value of ≤ 0.05. **D)** Abundance of tRNA species grouped according to their type of base interaction (i. Watson–Crick, ii. Watson–Crick and wobble (G:U), and iii. Watson–Crick and wobble (inosine) with their corresponding codons. Statistic significant tests (p value ≤ 0.05) were performed with the Wilcoxon–Mann‒Whitney test (for p values: * = 0.05; ns = not significant). E) Ratio of posttranscriptional modification signatures differently expressed (fold change log2≤ 0.25 between modification signatures) in the tRNA nucleotides positions from the EPI and MT forms. Percentage of 3’CCA, 3’CC or 3’C sequences at tRNA molecules in the EPI and MT forms. A Q-test was performed, and a p value of ≤ 0.01 was considered statistically significant for tRNA maturation and misincorporation ratios across different life forms of *T. cruzi*.

EPIs and MTs express different sets of genes. In particular, EPIs mainly express genes encoded in the core compartment, whereas MTs exhibit increased expression of genes encoded in the disruptive compartment (de Godoy et al., 2012; Smircich et al., 2015). Thus, we hypothesized that the tRNA availability in EPIs and MTs fit the codon usage of the core and disruptive compartments, respectively. We detected greater expression of tRNA Glu1B (CTC), Glu1A (CTC), and Thr2 (CGT) in EPIs than in MTs (p-adjusted <0.05) (Figure 3C and Supplementary Table 2D), which corresponds to two (Glu (GAG) and Thr (ACG)) of the five optimal codons in the core compartment (Figure 1G). In contrast, tRNAs corresponding to any of the five optimal codons from the disruptive compartment were not increased in MTs (Figure 1G). These data suggest that there is some, albeit limited, correspondence in the tRNA expression profile in EPIs to fit the core compartment, but the expression profile in MTs does not fit the disruptive compartment. Thus, tRNA abundance undergoes only minor modulation during the differentiation process from EPIs to MTs, despite significant differences in protein expression observed between these forms.

Furthermore, we found that tRNAs with anticodons exhibiting wobble base pairing are not more abundant than those relying solely on Watson–Crick base pairing. This suggests the lack of a regulatory mechanism to increase the abundance of wobble tRNAs to meet the demand for multiple codons (Figure 3D and Supplementary Table 2E).

In addition, since some tRNA modification causes misincorporation of wrong bases during RT, misincorporation frequency can be used to probe the frequency of tRNA modification. Thus, although the abundances of tRNAs in EPIs and MTs are similar, we identified a marked reduction of putative post-transcriptional modifications (Figure 3E and Supplementary Figure 2) and maturation in several tRNAs in MT compared to EPI by analyzing misincorporation and 3’CCA rates, respectively (Figure 3F). CCA end is a prerequisite for proper tRNA aminoacylation and therefore, to produce competent tRNAs. Thus, lower levels of o mature tRNAs in MT together with changes of modification states in two life forms can modulate the tRNA functions.

### The chromatin profile of tDNAs correlates with the abundance of their transcripts in the epimastigote form.

The tDNA copy number exhibits a strong positive correlation with transcript abundance in some eukaryotes (Behrens et al., 2021). Notably, we found only a weak correlation (Spearman’s *r* = ≈ 0.36) between tRNA transcript abundance and copy number in both the EPI and MT forms, indicating that the tRNA abundance profiles in *T. cruzi* are not solely defined by gene copy numbers (Figure 4A).

**Figure 4.**
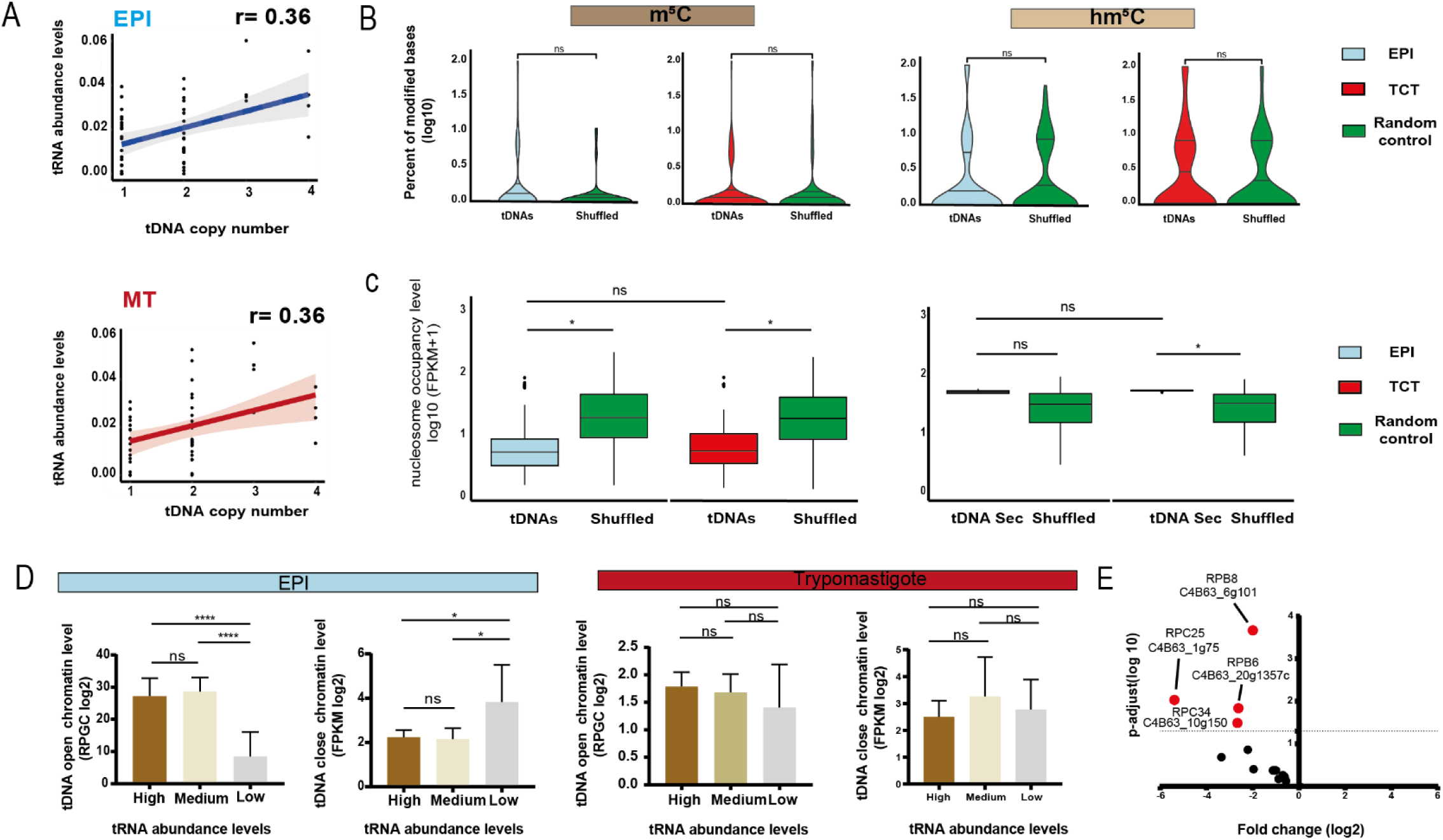
Correlation between tDNA chromatin and tRNA abundance in *T. cruzi* life forms. **A)** Spearman’s correlation between tRNA abundance and the number of corresponding tDNA copies in the EPI and MT forms. tRNA^Sec^ is not included in the scatter plot. Red or blue shading, 95% confidence interval. **B)** Percentage of m^5^C and hm^5^C modifications on tDNA nucleotides present in the EPI and TCT forms. **C)** Nucleosome enrichment in tDNAs and random controls (shuffled) in the EPI and TCT forms using fragments per kilobase per million mapped reads (FPKM) +1 (log10) values from MNase-seq data. Shuffled was used as random genomic controls. **D)** Bar chart showing the associations between high-, medium- or low-abundance tRNAs and their corresponding tDNA chromatin profiles. The open chromatin levels were normalized to the reads per genomic content (RPGC) from FAIRE-seq data obtained with the EPI and MT forms. The closed chromatin were normalized to the FPKM values from MNAse-seq data obtained with the EPI and TCT forms. Statistical significance tests were performed with the Wilcoxon–Mann‒Whitney test (for p values: **** = 0.0001; * = 0.05; ns = not significant). **E)** Volcano plot depicting the fold changes and p values of RNAP III subunits in *T. cruzi* life forms (EPI vs. MT) using Ribo-seq data and DESeq2 algorithm. A p value of ≤ 0.05 and fold change ≥ 1.5 were considered statistically significant.

We demonstrated that tDNAs from EPI forms exhibited greater levels of open chromatin than did those from their infective forms (Lima et al., 2022), suggesting a chromatin regulatory role. Here, we investigated whether tDNA chromatin profiles are associated with tRNA abundance. To this end, we used Nanopore, FAIRE-seq and MNase-seq datasets from different *T. cruzi* life forms (Díaz-Viraqué et al., 2023; Lima et al., 2022, 2021). The Nanopore datasets included information on DNA modifications, such as 5-methylcytosine (m^5^C) and 5-hydroxymethylcytosine (hm^5^C), which are associated with increased and decreased gene expression, respectively (Hummel et al., 2020; Shi et al., 2017). We found that tDNAs are not localized in regions enriched in m^5^C and hm^5^C modifications compared to the shuffled control (Figure 4B). In addition, the abundance of these DNA modifications was not associated with different tRNA expression levels in either the EPI or infective form (Supplementary Figure S2B).

MNase-seq assesses DNA regions associated with nucleosomes, providing information on nucleosome occupancy and positioning at any given locus (Chereji et al., 2019). We observed lower nucleosome occupancy levels in all canonical tDNAs in both the EPI and infective forms (TCTs), indicating that tDNAs tend to be localized in nucleosome-depleted regions (NDRs) (Figure 4C). However, tDNA loci encoding tRNA^Sec^ showed a different trend. The levels of nucleosome occupancy at the eleven tDNAs^Sec^ did not differ from those of the shuffled control, suggesting that the tDNA Sec loci are localized in nucleosome-enriched regions or closed chromatin.

Next, we used the FAIRE-seq dataset, which detects open chromatin, also known as transcriptionally active chromatin regions (Giresi et al., 2007; Simon et al., 2012). We obtained the abundance values of the twenty single-copy tRNA species for comparisons between their expression levels and chromatin status, mitigating the bias likely introduced by multicopy tRNA genes. The tRNAs were classified into three groups (high, medium- and low-abundance) based on the tRNA abundance obtained by tRNA-seq (Supplementary Figure S2A). We found that tDNAs with high and medium abundances exhibit more open chromatin compared to those with low expression in the EPI forms, suggesting that chromatin status serves as an important regulator of tRNA expression levels. However, the open chromatin profiles of tDNA did not exhibit a distinct pattern based on the levels of tRNA abundance in *T. cruzi* infective forms, suggesting the presence of different mechanisms regulating tRNA abundance in the infective form of *T. cruzi* (Figure 4D and Supplementary Table 3). Evaluation of nucleosome occupancy levels (MNAse-seq data) retrieved similar conclusions (Figure 4D). Notably, there is no current evidence that tRNAs are transcribed in nonproliferative/infective forms. To gain further insight into this issue, we evaluated the expression of the subunits of RNAP III, which are dedicated to tRNA transcription, in the EPI and MT forms. We detected a downregulation of multiple components of RNAP III in the MT form compared to the EPI form (Figure 4E and Supplementary Table 2F). This observation suggests a lower tRNA transcription activity in *T. cruzi* infective forms, which could account for the similar profile levels of tRNAs from EPIs to MTs.

### tRNA abundance coadapts with codon frequencies of highly translated transcripts

We investigated whether tRNA abundance exhibits coadaptation with the corresponding codon frequency. We found a moderate correlation (R=0.63) between codon frequency for all CDSs and tRNA abundance in EPIs (Figure 5A). Notably, core CDSs (R=0.63) exhibited a stronger correlation than did disruptive CDSs (R=0.42). As MTs express more disruptive proteins than EPIs (de Godoy et al., 2012; Freire-de-Lima et al., 2015; Smircich et al., 2015), we asked whether codon frequency and tRNA abundance are more strongly correlated for disruptive genes in MTs. However, similar to that for EPIs, the correlation between codon frequency for disruptive genes and tRNA abundance was lower than that for core genes (R= 0.33 and R=0.56) (Figure 5A), which is consistent with the data shown in Fig. 3C, where tRNA abundance profiles only slightly changed between EPIs and MTs.

**Figure 5.**
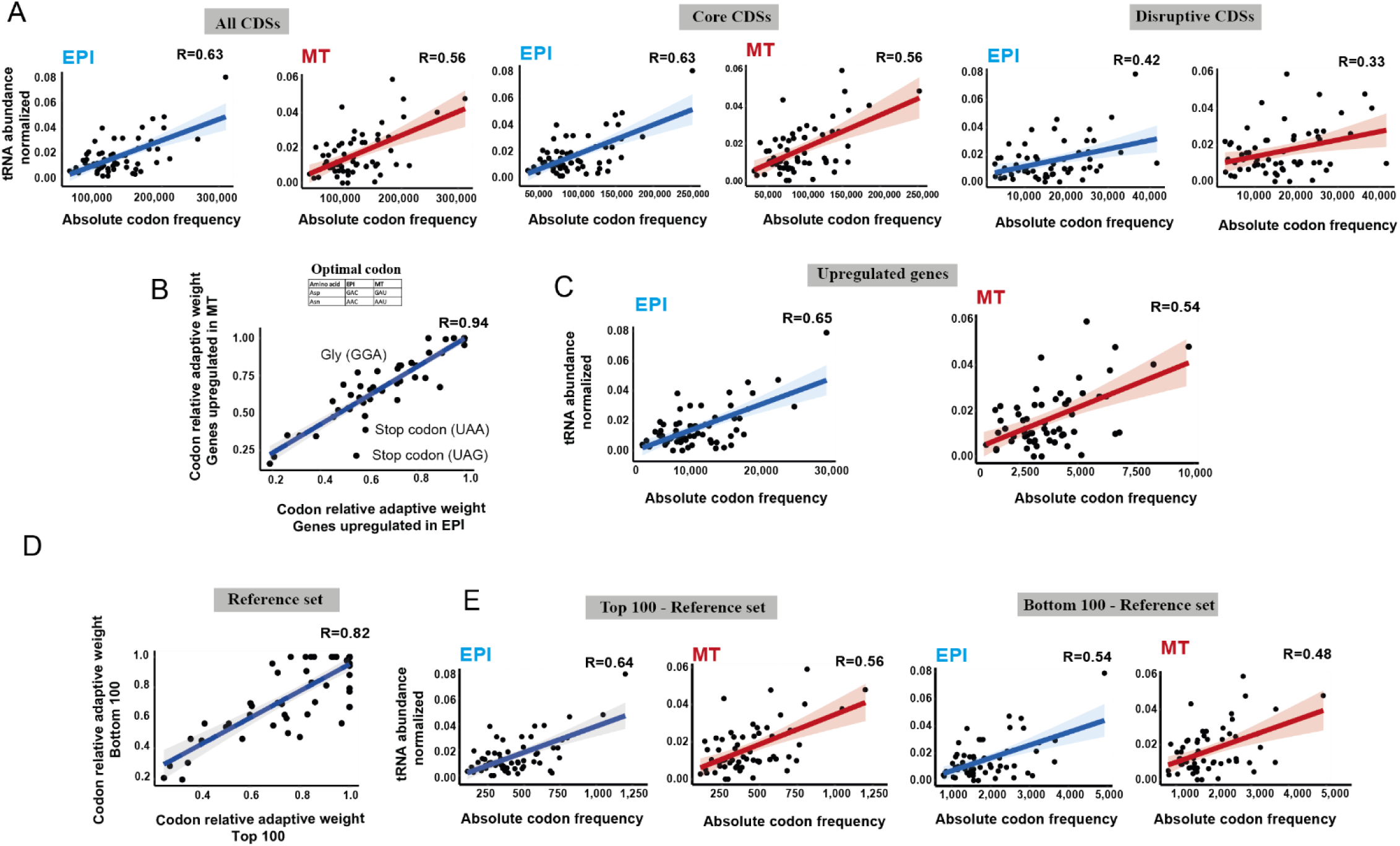
Relationship between tRNA abundance and codon usage frequency in *T. cruzi*. **A)** Pearson’s correlation between tRNA abundance in the EPI and MT forms with codon usage frequencies in all CDSs, core genes and disruptive genes. **B)** Pearson’s correlation for the relative adaptive weights of codons in the upregulated CDSs for the EPI (1,752 genes) and MT forms (361 genes), obtained with translatome data (Ribo-seq). The distinct optimal codons in the upregulated genes in the EPI and MT forms are illustrated in the table below the scatter plot. The highlighted codons are at least 20% different between the indicated conditions. **C)** Pearson’s correlation between tRNA abundance in the EPI and MT forms with their absolute codon frequencies from upregulated genes. **D)** Pearson’s correlation for the relative adaptive weights of codons in the top 100 and bottom 100 genes from a reference dataset containing more and less abundant transcripts translated from *T. cruzi* life forms. **E)** Pearson’s correlation between tRNA abundance in the EPI and MT forms with codon frequencies of higher (top 100) and lower (bottom 100) translated transcripts. The expression level was based on transcripts per million (TPM) values from Ribo-seq data. Shading-95% confidence interval.

Next, we asked whether the codon frequency of highly expressed transcripts in EPIs or MTs matches their tRNA pools. The genes upregulated in the EPIs compared to the MTs (Figure 5C and Supplementary Table 1B-E) had a similar correlation with all of the other CDSs (R=0.65 vs. R=0.63) (Figure 5A and C). The genes upregulated in MTs compared to EPIs (Figure 5C) also exhibited a similar correlation to all CDSs (R=0.54 vs. R=0.56) (Figure 5A and Supplementary Table 1B-E), indicating that differentially expressed genes do not have a more optimal codon bias for efficient expression in either form.

Subsequently, we evaluated the association between codon frequency and tRNA abundance for the more abundant and less abundant translated transcripts from the reference set (Supplementary Table 1D-E), which were created to avoid multiple-copy genes and transcripts that exhibited large differences (fold change log2 ≥ 2) in expression between life forms. We found that the correlation was greater for the highly abundant transcripts (top 100) than for the less abundant transcripts (bottom 100) in both EPIs and MTs, suggesting that a regulatory mechanism based on tRNA availability operates similarly across *T. cruzi* life forms (Figure 5E). However, the correlations between tRNA abundance and the corresponding codon frequencies in MTs were lower than those observed in EPIs.

Finally, we compared the codon usage bias (considering the codon adaptive weights—both the optimal and nonoptimal codons) among all groups of genes mentioned above. We found that codon bias was most pronounced when comparing the core and disruptive compartments (R=0.75), followed by the top 100 and bottom 100 genes (R=0.82) and finally the upregulated genes (R=0.94) (Figure 1H, Figure 5B and 5D).

### Base pairing modes between codons and anticodons are linked to translated transcript levels

To further understand the observation that highly expressed transcripts (top 100) are more strongly correlated with tRNA abundance than are less expressed transcripts (bottom 100) (Figure 5E), we performed a detailed analysis of the codon composition of these genes. We identified forty-six tRNA anticodons and sixty-two tRNA codons in the *T. cruzi* genome, implying that some tRNA transcripts must recognize more than one codon via tRNA modification (inosine) or wobble base pairing (G-U) (Figure 2D). We found nine distinct optimal codons between the top 100 or bottom 100 transcripts (Figure 6A). With the exception of Ala (GCC/GCG), all of these residues are recognized by the same set of tRNA species. Strikingly, six (out of nine) optimal codons from the top 100 abundant transcripts interact via Watson–Crick base pairing only, while six (out of nine) optimal codons from the bottom 100 form only wobble base pairing (Figure 6B and C). This observation suggested that the anticodon:codon base pairing mode is associated with the expression level of the transcripts in *T. cruzi*.

**Figure 6.**
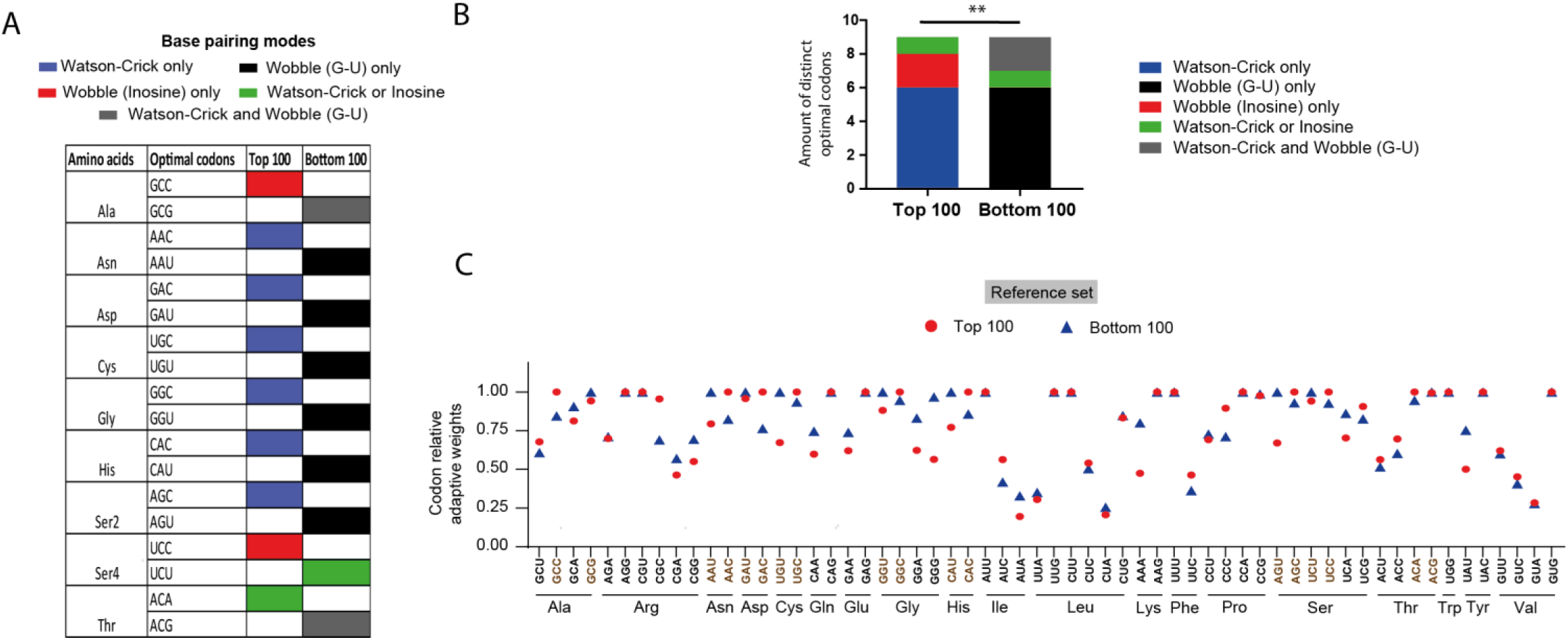
Optimal codons and anticodon:codon pairing modes in the top 100 and bottom 100 genes in *T. cruzi*. **A)** Distinct optimal codons between the highest 100 (top 100) and lowest 100 (bottom 100) translated transcripts in the EPI and MT forms. **B)** Distribution of the anticodon:codon base pairing modes (**i.**Watson–Crick only, **ii.** wobble (G-U) only, **iii.** wobble (inosine) only, **vi.** Watson–Crick and wobble (inosine) and Watson–Crick and wobble (G:U)) were present in distinct optimal codons between the top 100 and bottom 100 genes. Chi-square - p value ≤ 0.01 (**) was considered. **C)** Codon relative adaptive weights compared to optimal codons from the top 100 and bottom 100 genes.

## Discussion

Codon usage bias has been proposed to be a significant factor directing transcript and protein expression levels in trypanosomes (de Freitas Nascimento et al., 2018; Jeacock et al., 2018). Codon frequency is linked to the abundance of tRNA pools in some species, but measuring tRNA expression levels has been only a recent advancement (Behrens et al., 2021). Here, we sequenced tRNAs in two different developmental forms of *T. cruzi*, which enabled direct comparisons between tRNA abundance and codon usage biases. We observed a stronger correlation between codon usage and tRNA abundance than between the copy number of tRNA genes (Figure 4A), highlighting the importance of obtaining actual tRNA expression profiles to analyze the relationship between codon bias and tRNA supply. Surprisingly, tRNA abundance profiles were very similar in EPIs and MTs (Figure 3C), although these two forms have distinct gene expression profiles (de Godoy et al., 2012; Smircich et al., 2015). tRNA abundances correlated with the core compartment of the genome and highly expressed mRNA transcripts in both life forms (Figure 5A, C and E).

The transcription of tRNAs relies on RNA Pol III and interactions between transcription factors and promoters within tRNA loci. This process is facilitated by the depletion of nucleosomes at these loci (Graczyk et al., 2018; Shukla and Bhargava, 2018). We showed that tDNAs are in regions that are lacking in nucleosomes in both the EPI and TCT forms, except for tDNA^Sec^ (Figure 4C). This exception could be explained by the fact that tRNA^Sec^ is transcribed by RNA Pol II, unlike canonical tRNAs (Aeby et al., 2010). Consequently, these results indicate that the chromatin status can influence tRNA transcription (Figure 4D). Notably, this correlation was exclusive to EPI forms (Figure 4D). In trypomastigote forms (both TCTs and MTs), tRNA expression appears to be independent of chromatin status, suggesting that other regulatory mechanisms maintain the tRNA abundance profiles in MTs similar to those in the EPI form. Considering the presence of NDRs in the tDNA of MTs (as observed in TCT forms), it is plausible that chromatin closure in MTs (as observed in FAIRE-seq assays) (Lima et al., 2022) may be attributable to the deposition of proteins associated with chromatin compaction but due to nucleosome deposition. Future investigations are warranted to validate this hypothesis.

Another possible explanation for the similar tRNA abundance profiles in the MT and EPI forms is that de novo tRNA synthesis is not active in the MT form, resulting in maintenance of the same tRNA profiles as in the EPI form. The low amount of mature tRNAs (per parasite) in MT forms (Lima et al., 2022), alongside a downregulation of the RNAP III machinery in these forms (Figure 4E), suggests that tRNA synthesis is not active in MTs, while nascent expression via RNA Pol III has not yet been assessed in *T. cruzi*. Pol III RNA expression can also be arrested by Maf1, which was described in *T. brucei* to inhibit RNAP III transcription in procyclic forms (Romero-Meza et al., 2017). Since Maf1 is known to bind to RNAP III under stress conditions in some organisms (Graczyk et al., 2018), we hypothesized that a similar mechanism may also play a role during *T. cruzi* differentiation.

To assess the effect of codon bias on *T. cruzi* gene expression, we evaluated genes from the core and disruptive regions and genes with higher and lower abundances. Genes in the disruptive regions showed increased expression in *T. cruzi*’s infective forms (de Godoy et al., 2012; Smircich et al., 2015) but comprised only three gene classes, while the core compartment encompassed all of the remaining CDS (Berná et al., 2018). Therefore, we expected these two compartments to differ greatly in codon usage, which was confirmed in the present study (Figure 1H). Additionally, we also found nine pairs of codons that were optimized differently among the most and least abundant transcripts (Figure 6A). Together, these results indicate the presence of codon bias associated with both gene expression and genomic compartments in *T. cruzi* life forms. Notably, we found that the most and least abundant transcripts use codons that are more or less adapted to their tRNA pool (Figure 5E), suggesting that the tRNA abundance profile is adapted to codon usage in highly expressed genes.

We also found that the stop codon UGA is prevalent in disruptive genes, while the core and GpDR compartments contain a homogeneous distribution of the three stop codons present in the *T. cruzi* genome (Figure 1E). The enrichment of the UGA stop codon in disruptive genes could confer a significant advantage, as UGA is the least efficient stop codon compared to UAA and UAG in eukaryotes, thereby increasing the likelihood of translation readthrough (Anzalone et al., 2019; Belin and Puigbò, 2022; Poole et al., 1995; Trexler et al., 2023). This phenomenon could potentially allow for the translation of additional C-terminal sequences and domains, enhancing protein variability that may be crucial for these antigenic groups of proteins (Schueren and Thoms, 2016). In addition, the UGA codon can be recognized by tRNA^Sec^ if a SECIS element is present downstream of the mRNA (Berry et al., 1993; Mariotti et al., 2013). Interestingly, we found a fourfold increase in the level of tRNA-Sec in MTs (nonproliferative cells) (Figure 3C). Among the disruptive genes enriched in UGA, we found that only 1.4% of genes (mainly TSs) had putative SECIS elements (data not shown), indicating that in the majority of cases, UGA at disruptive genes is indeed associated with a stop in translation and not with the insertion of a selenocysteine amino acid.

Our results suggest that tRNA abundance may be less coadapted to gene expression in MTs than in EPIs. However, this coadaptation can also be further modulated by the presence of PTMs in tRNAs. In this regard, we found that tRNAs from MTs contain fewer putative PTMs, mainly those located within the tRNA body (Figure 3E), which can affect stability (Cui et al., 2023; Ohira and Suzuki, 2024; Yared et al., 2024). Thus, it is plausible that in MTs, the coadaptation of tRNAs and codon frequency may be even lower. Evaluating the full repertoire of PTMs in tRNAs of different life forms will shed light on the role of these PTMs in tRNA biology and gene expression in *T. cruzi*.

Trypanosomatids have fewer copies of tRNA genes (fewer isodecoders) (Figure 2A) than vertebrates (Goodenbour and Pan, 2006; Santos and Del-Bem, 2022), suggesting that these genes may be under selective pressure to avoid deleterious mutations. Here, we detected forty-six anticodons for sixty-two codons (Figure 2D and Supplementary Table 2B), which contrasts with findings in humans, where there are fifty-seven anticodons for sixty-one codons (Iben and Maraia, 2014; Kanduc, 2021). This indicates that the parasite needs to employ more strategies to recognize the entire codon repertoire of the genome, such as employing more modifications in anticodons and codon‒anticodon wobble pairing. In this sense, we found that *T. cruzi* may take advantage of another layer of gene expression regulation based on anticodon:codon pairing modes. We found that the most abundant genes from each life form predominantly use codons that utilize Watson–Crick base pairing (Figure 6A–C). In contrast, the least abundant transcripts had greater frequencies of wobble-pairing codon‒ anticodon interactions. tRNA pairing affects translation efficiency, with the latter type being associated with lower efficiency (Chan et al., 2017; Stadler and Fire, 2011). Preferential usage of codons forming Watson-Crick pairing in highly expressed transcripts were also observed in other organism, such as *Plasmodium falciparum* (Chan et al., 2017). Such skewed synonymous codon usage can be a common strategy to optimize translation among parasites.

Future studies should experimentally assess whether the expression of a protein is indeed affected by swapping codons recognized by different pairing modes. In brief, we propose that *T. cruzi* protein expression in EPIs and MTs may be affected by a combination of codon usage bias, tRNA abundance and anticodon:codon pairing mode. Subsequent studies should explore the potential impact of perturbed tRNA biology on parasite viability, thereby offering promising avenues for novel disease-control strategies, particularly in the realm of tRNA-based therapeutics.

## Supplementary information

**Figure S1.**
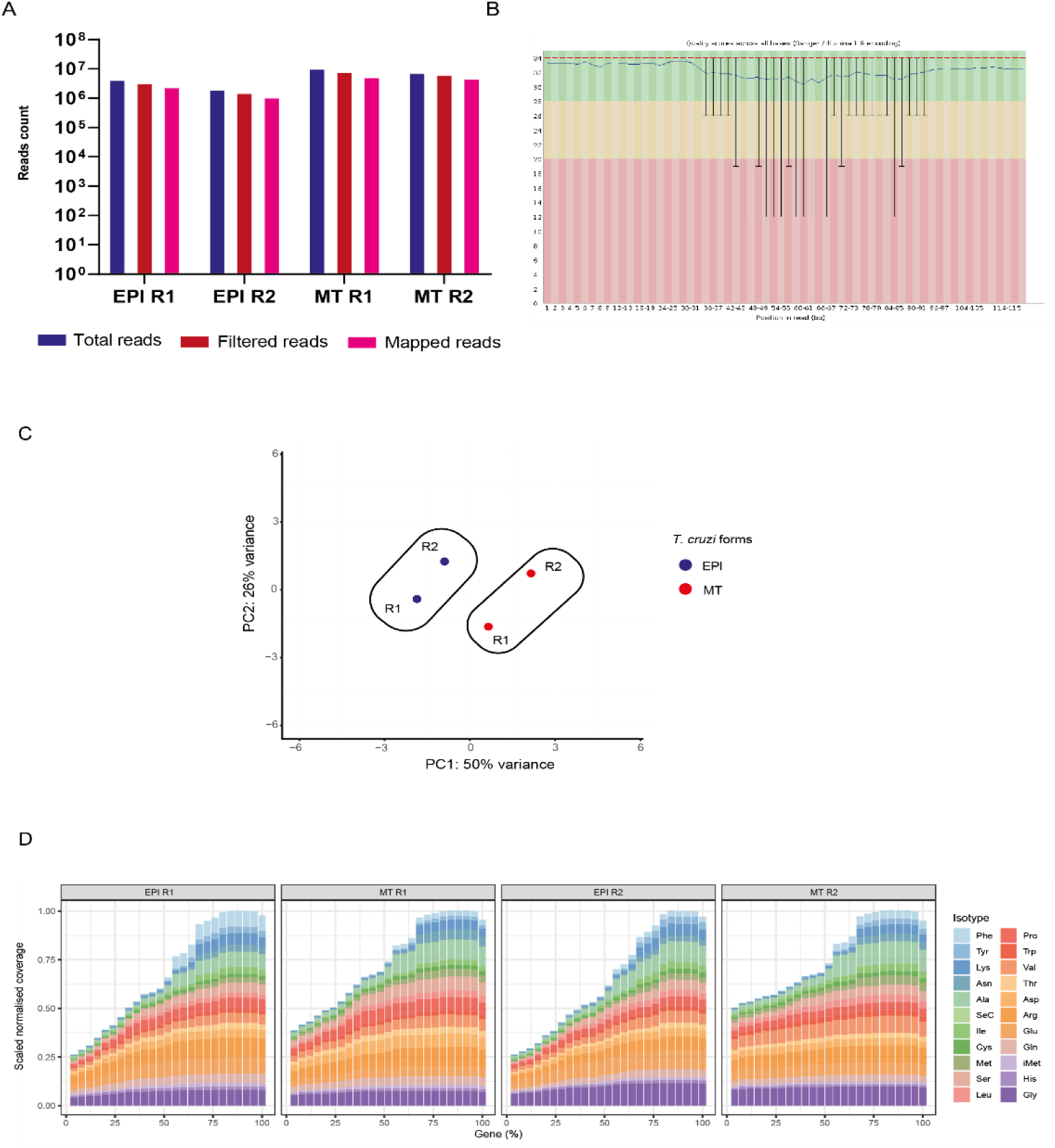
Quantity of reads and sequencing data quality for tRNA-seq performed in the EPI and MT forms. **A)** Number of reads sequenced, filtered and mapped on tDNAs from *T. cruzi* Dm28c. **B)** Quality of base pair sequences, assessed by Phred scores, of a representative sample (MT R1). **C)** PCA plot of tRNA-seq-normalized samples with the varianceStabilizingTransformation tool from the DESeq2 package. R1= replicate 1; R2= replicate 2. **D)** Metagene analysis of the normalized sequence coverage of tRNA isotypes.

**Figure S2.**
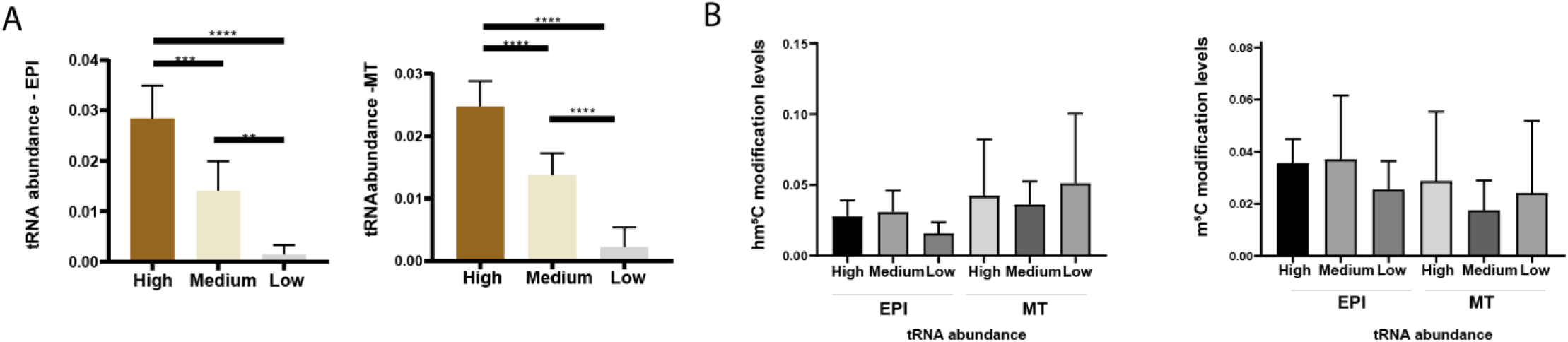
Abundance of twenty tRNA transcripts of a single copy in *T. cruzi.* **A)** Bar charts depicting the seven tRNAs with the highest abundance, the seven with medium expression, and the remaining six with the lowest abundance in either the EPI or MT form. **B)** Relationships between tRNA abundance levels (high, medium and low) and hm^5^C or m^5^C modification levels in twenty single-copy tDNAs in the EPI and MT forms. Statistical significance tests were performed with the Wilcoxon‒Mann‒Whitney test. (ns = not significant).

## ADDITIONAL FILES

**Table 1.** Codon usage in the *T. cruzi* genome. **A)** Codon frequencies, RSCU and codon relative adaptive weight in the core, disruptive and GpDR compartments. **B)** TPM values for all genes in the EPI and MT forms obtained from the translatome data (Ribo-seq). **C)** DESeq2 results from Ribo-seq data performed in the EPI and MT forms. **D)** Gene IDs of the reference set (top 100 and bottom 100) and genes upregulated in the EPI and MT forms. E) Analysis of the distribution of optimal codons, codon frequencies, and codon adaptive relative weight for either the top 100 or bottom 100 genes in a reference set, as well as for the dataset of genes upregulated in the EPI and MT forms.

**Table 2.** Characterization of tDNA and tRNA pools in *T. cruzi* Dm28c. **A)** Genomic coordinates of tDNAs, gene IDs, sense strands and copy numbers. **B)** Identification of tRNA anticodons and their base pairing types with corresponding codons in the *T. cruzi* genome**. C)** tRNA abundance of replicates (R1 and R2) and their averages in both the EPI and MT forms. **D)** DESeq results of tRNA abundance in EPIs compared to the MTs. **E)** Correlations between tRNA abundance and base pairing types with their corresponding codons. **F)** BLAST and HMMER results containing the best hit of the RNAP III subunit candidates present in the *T. cruzi* genome.

**Table 3.** Abundance of single-copy tRNAs and their levels of open (FAIRE-seq) and closed (MNase-seq) chromatin in the EPI and MT forms.

## Author Contributions

Silva, H.G. and Cunha, J.P.C. contributed to the writing, analysis, and rationale of this article. Kimura, S. and Waldor, K. M. assisted in executing the tRNA sequencing. Lima, P.L.C. obtained the RPKM values from the MNase-seq data, and Pires, D obtained the tDNA modification data. All authors contributed to the revision of this work.

## Acknowledgments

We thank Ivan Novaski Avino and Karin Navarro for technical assistance and Dr. Alex Ranieri Lima for important input on the data and bioinformatic analysis. We thank Dr. Janaina de Freitas Nascimento and Dr. Mark Carrington for their valuable discussions on codon usage data. We thank Leticia de Sousa Lopes for reviewing this manuscript.

## Data availability

The t-RNA-seq, MNAse-seq, and FAIRE-seq data can be accessed at the Sequence Read Archive (SRA) (https://www.ncbi.nlm.nih.gov/sra) under the following accession numbers: PRJNA1124437, PRJNA665060 and PRJNA763084. Oxford Nanopore sequencing datasets are available at(28).

