## Supplementary figures and images for "Integrating tDNA Epigenomics and Expression with Codon Usage Unravel an Intricate Connection with Protein Expression Dynamics in *Trypanosoma cruzi*"

### Figure S1

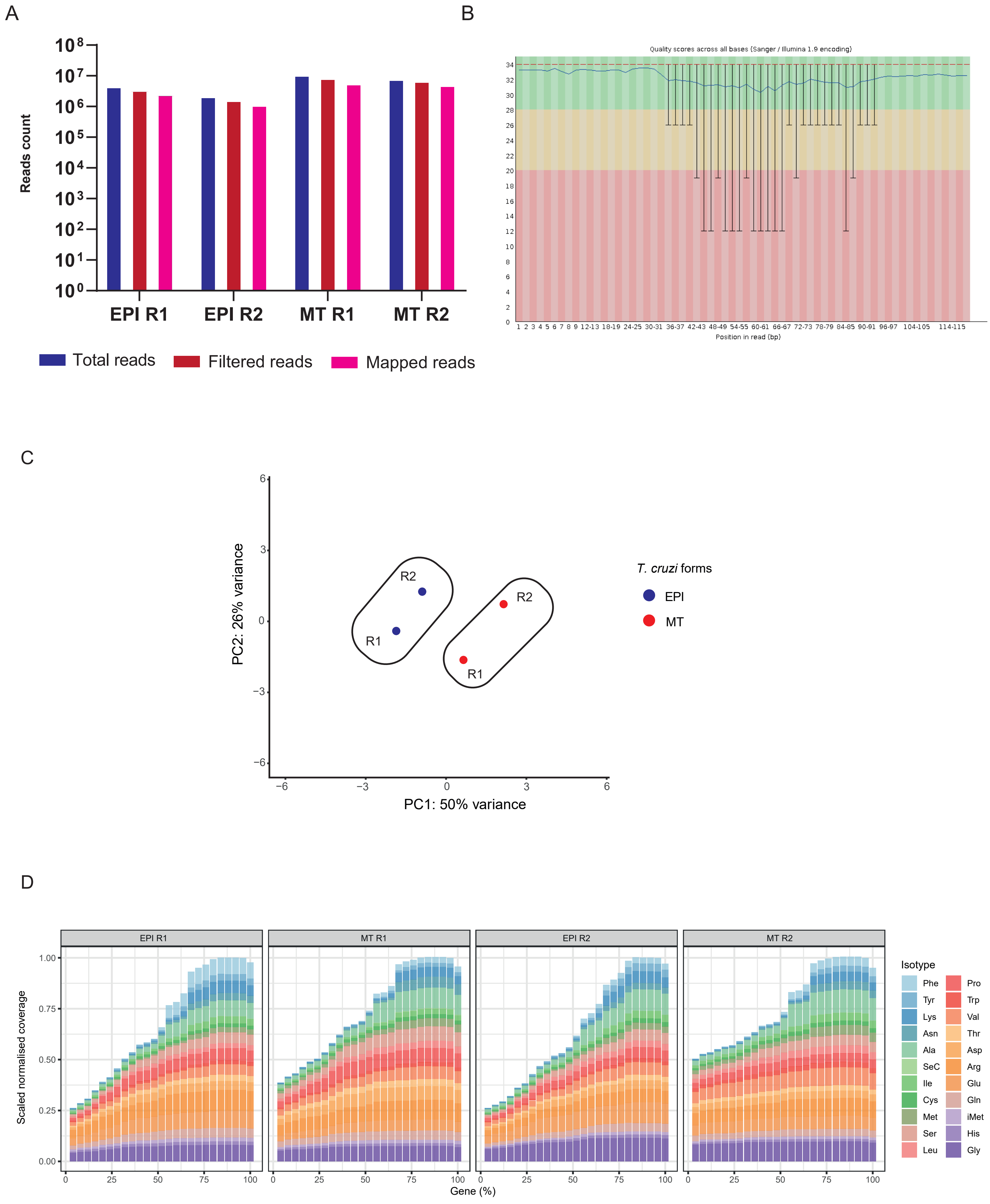

### Figure S2

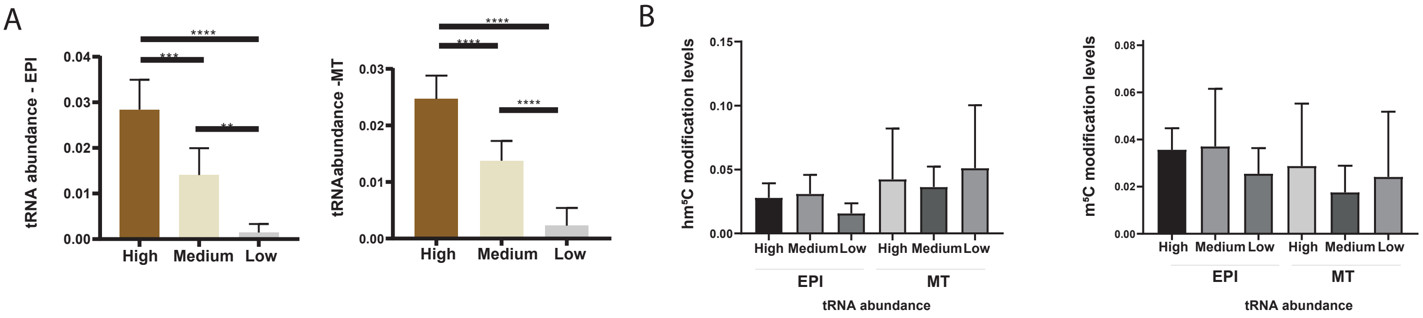
